# Adult *Drosophila* legs do not regenerate after amputation

**DOI:** 10.1101/2022.10.25.513553

**Authors:** Anne Sustar, John C. Tuthill

## Abstract

A recent paper by Abrams *et al*. (2021) claimed that a simple dietary supplement is sufficient to induce appendage regeneration in jellyfish, flies, and mice. This would be remarkable, if true, because it was previously thought that flies and mice lack the capacity for regeneration after injury. We therefore sought to replicate their provocative results. We amputated one tibia of over 1000 fruit flies, fed them control or supplemented diets, and carefully examined their legs three weeks post-injury. We did not, however, observe any instances of leg regeneration. We conducted additional experiments that confirmed the complete absence of neurons, muscles, or other living cells in amputated tibias. Abrams *et al*. also reported the formation of a white blob at the amputation site, which they interpreted as an intermediate regeneration morphology. We tested this hypothesis more rigorously and conclude that the white blob consists of bacteria. Overall, we failed to find any evidence for leg regeneration in *Drosophila*, even when flies were fed the supplemented diet. Our results therefore contradict the overarching conclusion of Abrams *et al*. that dietary supplements are sufficient to unlock an ancestral mechanism that induces appendage regeneration.

## Introduction

Abrams *et al*. (2021) reported that supplementing an animal’s diet with L-leucine and insulin/sucrose promotes appendage regeneration, even in species that were previously thought to lack the capacity for regeneration. The potential discovery of a universal means to unlock regenerative capacity is exciting because it could be applied to other animals, including humans. Indeed, a central conclusion of Abrams *et al*. is that their “study suggests that an inherent ability for appendage regeneration is retained in non-regenerating animals and can be unlocked with a conserved strategy.”

We initially became interested in the specific part of the Abrams *et al*. study that addressed limb regeneration in the fruit fly, *Drosophila melanogaster*. Because we study proprioception and motor control of the *Drosophila* leg, we were curious to investigate how neurons and muscles regenerate in injured limbs. We initially focused on replicating the finding that fruit flies can, rarely and under specific experimental conditions, regenerate amputated legs.

Appendage regeneration has been extensively studied in insects. Some groups of insects have been shown to regenerate whole limbs, while others not at all. In general, regeneration capacity is linked to how an insect develops through metamorphosis (Yang et al., 2016). Hemimetabolous insects, such as stick insects, cockroaches, and crickets, have incomplete metamorphosis – they develop as nymphs that resemble small adults. Nymphs possess miniature versions of adult appendages, including legs, antennae, and in some cases, wings. As the nymph progresses through instars, developmental programs periodically break down and rebuild ectodermal tissues, depositing a new chitinous cuticle each time. So when the limb of a hemimetabolous insect is amputated, it can reactivate the same developmental patterning pathways and replace missing structures, including ectodermal tissue (Bando et al., 2018; Bodenstein, 1955; Bohn, 1971; French et al., 1976).

In contrast, holometabolous insects, such as flies, butterflies, and beetles, undergo complete metamorphosis. They initially develop into larvae in which the adult limb primordia are set aside as imaginal tissues. For example, cells that will become the adult legs are specified as distinct progenitor populations during embryogenesis. Later, during larval and/or pupal life, these cells undergo extensive proliferation, and ultimately differentiate and undergo morphogenesis to form the adult ectodermal structures during pupation. It is well known that during this early developmental phase, prior to differentiation, imaginal discs possess the capacity to regenerate following experimental removal of pieces of tissue (Fox et al., 2020; Haynie and Bryant, 1976; Schubiger, 1971). After metamorphosis, however, adult holometabolous insects never molt, and have never been found to regenerate lost or damaged appendages. Holometabolous insect limbs such as fly legs are thought to lack the developmental programs required to re-establish patterning and tissue growth after injury, likely due to epigenetic silencing of developmental genes (Fox et al., 2020; Harris et al., 2020; Repiso et al., 2011).

We began by repeating the fly leg amputation experiments of Abrams *et al*. We copied the protocol described in their paper, and through consultation with the senior author. We used the same wild-type *Drosophila* strain (Canton-S), experimental timeline, amputation site, and dietary conditions. Because Abrams *et al*. observed regeneration in only ∼1% of flies fed the supplemented diet, we used a sample size of ∼1000 flies, comparable to that described in their paper. When we failed to find evidence for cuticle regeneration, we next searched for specific tissues within the amputated leg stump to test whether flies are capable of regenerating neurons, muscle, or any other cell class. We also carefully examined the white blob at the cut site, a structure that Abrams *et al*. interpreted as an intermediate regeneration morphology.

Overall, we found no evidence for regeneration of any cells in the amputated leg. Rather, we find that upon amputation, the cells in the amputated leg segment all died and did not grow back. Furthermore, we provide evidence that the white blob growing on the amputation site is not a regeneration blastema, but rather a colony of bacteria. Our conclusions are consistent with the past literature supporting a lack of limb regeneration in *Drosophila* and other adult holometabolous insects.

## Results

### Absence of evidence for regeneration after amputation of *Drosophila* legs

Abrams *et al*. concluded that fly legs, which normally do not regenerate after amputation, show some regeneration ability when the fly’s diet is supplemented with insulin, leucine, and glutamine. We carefully followed their methods to replicate the fly leg regeneration experiments in Figure 3 of their paper. We amputated legs of 1283 flies, one hind leg per fly, at the midpoint of the tibia **(Figure 1; Table 1)**. The majority of these flies, 1083, were of the same wild-type fly strain (Canton-S) used in their study. After amputation, we raised 240 flies on control food and 843 on treated food. Three weeks later, we examined the legs at high magnification using bright-field microscopy, with the experimenter blind to experimental condition.

**Table 1.**
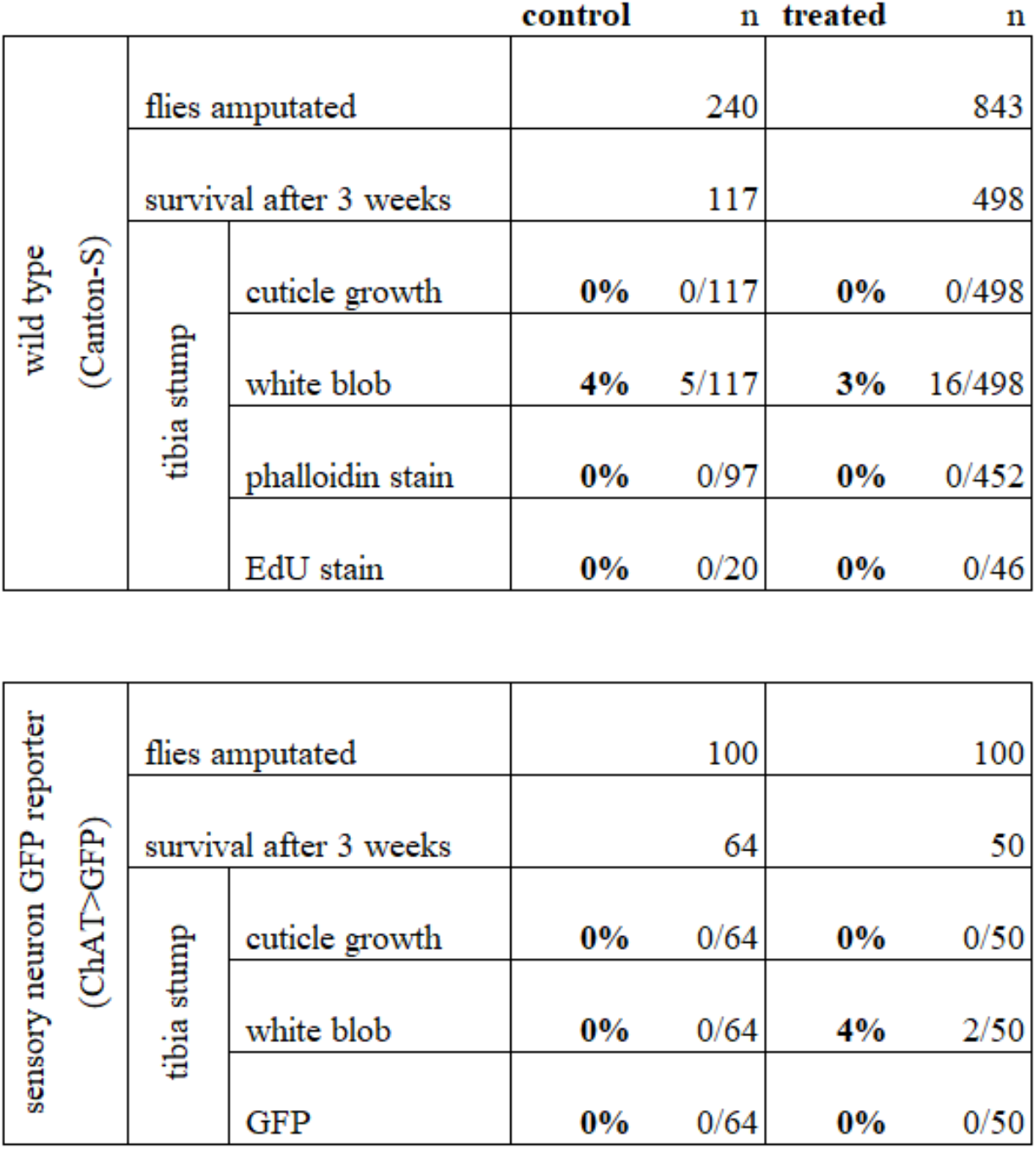
Summary of fly tibia amputation results.

We did not observe any regrown tibias, either in the control group or the treated group. The outcome of the two groups was qualitatively similar **(Figure 1)**. All tibia stumps had bristle deterioration and darkened cuticle, indicating necrosis. The site of the darkened cuticle varied. It was usually near the cut site, but sometimes farther up the leg, closer to the tibia-femur joint. In 4% of control cases and 3% of treated cases, we observed a white blob near the cut site **(Table 1; Figure 1, bottom row)**. The white blob was also observed by Abrams *et al*. to occur at a similar frequency. Overall, based on close inspection of the cuticle three weeks after amputation, we found no evidence of leg regeneration.

**Figure 1.**
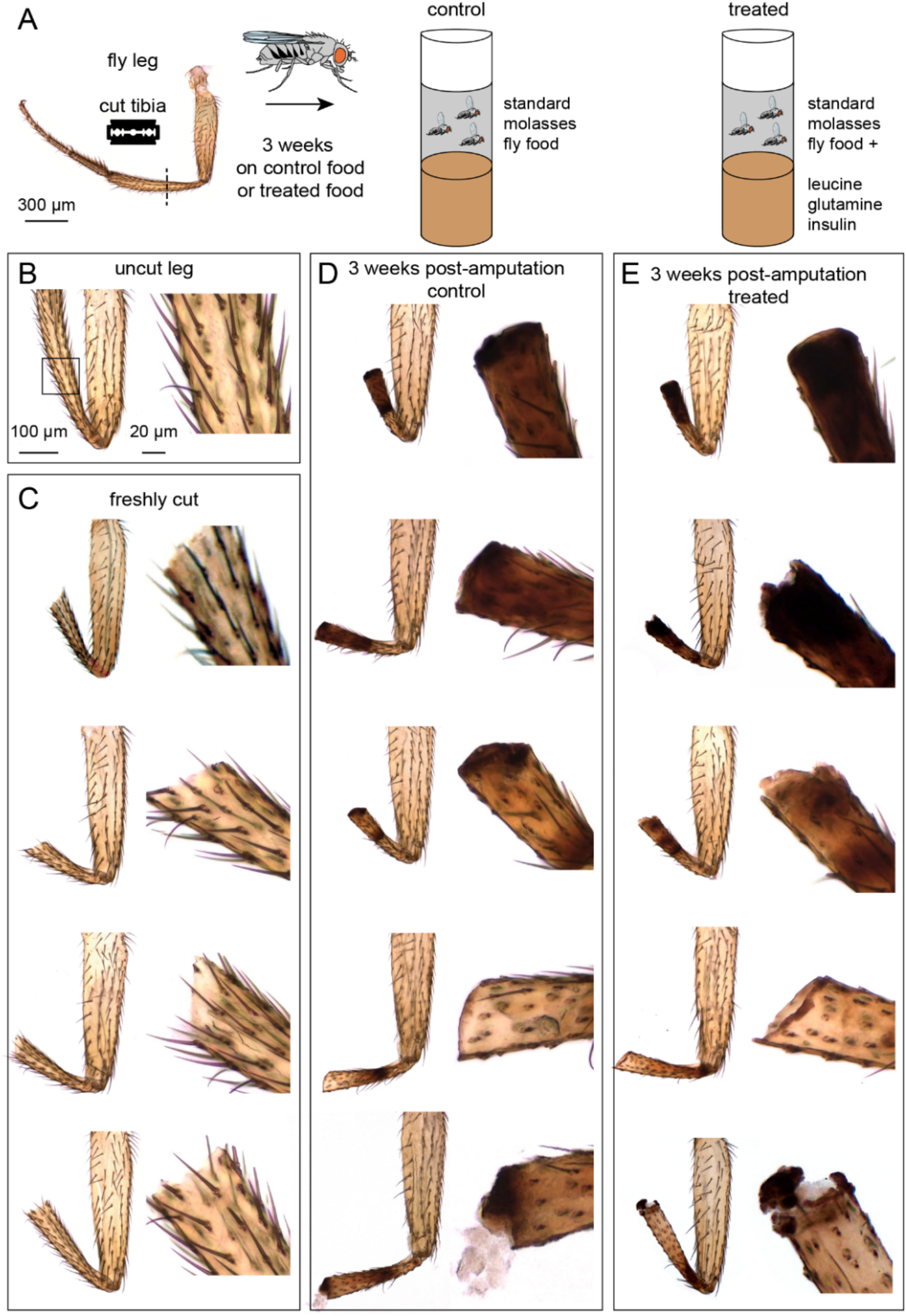
Tibia amputation in wild-type (Canton-S) *Drosophila* legs. (A) The experimental protocol. We amputated one hind leg per fly, at the midpoint of the tibia. After three weeks on control food or treated food, legs were fixed and analyzed. (B-E) Bright-field images of a control leg (B), four examples of freshly cut legs (C), and five examples of legs after three weeks on control food (D) or treated food (E). Insets showed magnified views of the cut site. Scale bar in B is the same for other panels.

### All cells die and fail to regenerate in the amputated tibia stump

Most of the internal structures of the fly leg, including muscle and neurons (Li et al., 2022), are largely transparent. Inspection of amputated limbs using bright-field microscopy alone might be insufficient to detect surviving or regenerated tissue. We therefore used fluorescent labels to test for the presence of muscles, neurons, and other cells in the tibia before and after amputation.

Each fly leg has ∼500 tactile bristles, including ∼120 distributed uniformly along the tibia (Schubiger and Hadorn, 1968). Each bristle is innervated by a single sensory neuron. To ask whether bristle sensory neurons can regenerate after injury, we amputated 200 legs in a fly strain with a fluorescent reporter that labels bristles and other sensory neurons (ChAT-Gal4>UAS-GFP). We raised half of the flies on control food and half on food with the supplemented diet. We then used confocal imaging to image GFP expression in the leg.

Although bristle neurons were present immediately after amputation, they were absent three weeks later, presumably due to degradation following cell death. The external bristle hairs also deteriorated and did not reappear. The results were indistinguishable between the control and treated groups **(Figure 2A; Table 1)**. In the femur and other leg segments proximal to the tibia, the bristle sensory neurons and hairs appeared normal. In summary, we failed to find any evidence that leg sensory neurons regenerate following tibia amputation.

**Figure 2.**
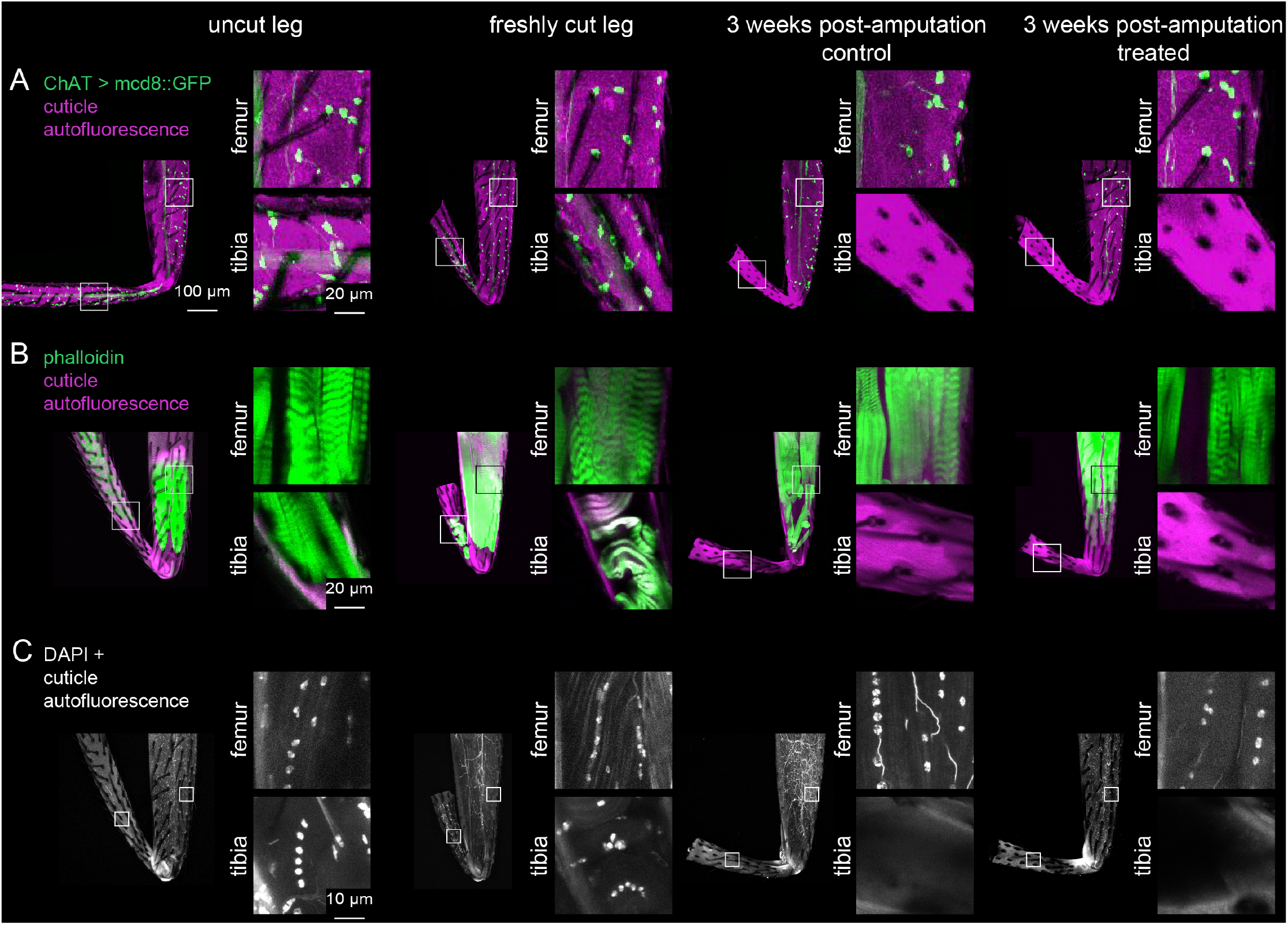
Investigation of tissue identity in amputated legs. Sensory neurons were labeled with ChAT-Gal4 > UAS-GFP (A), muscles were labeled with phalloidin (B), and nuclei were labeled with DAPI (C) in uncut legs, freshly cut legs, and 3 weeks post-amputation with control food or treated food. Insets show magnified views of tissue within the femur and tibia. Scale bars in A are the same for all other panels except for insets in C.

Each fly leg has twelve muscles, including four in the tibia (Soler, 2004). We next tested whether these tibia muscles can regenerate after amputation. We fluorescently labeled wild-type (Canton-S) leg muscles with phalloidin, a stain that labels F-actin. Three weeks after tibia amputation, neither the control group nor the treated group had any muscle staining in the tibia stump **(Figure 2B; Table 1)**. In the femur and other leg segments proximal to the amputation site, muscle staining was normal. In summary, we failed to find any evidence that leg muscles regenerate following tibia amputation. Since we did not find any sensory neurons or muscles in the amputated tibia stumps, we asked whether any other tissues, possibly hemocytes, glia, or epithelial cells, survive and/or regenerate. We stained legs of wild-type (Canton-S) flies with DAPI to label nuclei. Three weeks after amputation, we did not observe any DAPI staining in the cuticle of the tibia stumps, either in the control or treated groups (n=137 control, n=544 treated; **Figure 2C)**. In the femur and other leg segments proximal to the tibia, DAPI staining looked normal. This pattern is consistent with previous work showing that cell death was constrained to the injured leg segment in the adult cockroach (Bodenstein, 1955). In summary, our evidence supports the conclusion that all cells in the tibia stump die after amputation and fail to regenerate. The amputated stump appears to be an empty tube of cuticle, devoid of living cells.

### The white blob on amputated leg stumps is not a regeneration blastema

Abrams *et al*. reported the occasional appearance of a white blob at the tip of the amputated tibia stump in flies fed the supplemented diet. They interpreted this blob as an intermediate regeneration morphology. We observed the white blob form with a similar probability to that reported by Abrams *et al*. (3-4% of amputations, **Table 1**); however, we found that the blob occurred in both the control and experimental groups. Nonetheless, we sought to determine the nature of the white blob, and if it was, in fact, a sign of regeneration.

In many regeneration model systems, tissue regrowth is mediated by a blastema: a concentrated group of undifferentiated cells near the amputation site that proliferates to grow and repattern the missing tissue (Kiehle and Schubiger, 1985; Lehrberg and Gardiner, 2015). One possibility we considered is that the white blob at the cut site in amputated fly legs is a blastema. Alternatively, we thought that the blob could be fly tissue (e.g., muscle) extruded from the leg, a phenomenon which we sometimes observed immediately following amputation. To distinguish between these possibilities, we performed 5-ethynyl-2′-deoxyuridine (EdU) labeling on amputated legs (**Figure 3A-B)**. EdU is a thymidine analog that is incorporated into the DNA of proliferating cells during S-phase. It has been used extensively to label cell proliferation in regeneration blastemas of many animals, including *Drosophila* imaginal discs (Lehrberg and Gardiner, 2015; Worley et al., 2022). We amputated wild-type (Canton-S) fly legs, as above, and fed flies EdU continually for three weeks with either control food or the supplemented diet. When we harvested the legs for EdU staining, we also saved the fly gut as a positive control. Gut is one of the few adult fly tissue types that exhibits homeostatic cell proliferation during the adult stage (Xiang et al., 2017). All fly guts (n=5) had robust EdU staining **(Figure 3B)**. However, none of the tibia stumps had EdU staining **(Table 1)**, including legs with the white blob **(Figure 3A)**. We conclude that the amputated stump lacks regenerating cells and that the white blob is not a regeneration blastema.

**Figure 3.**
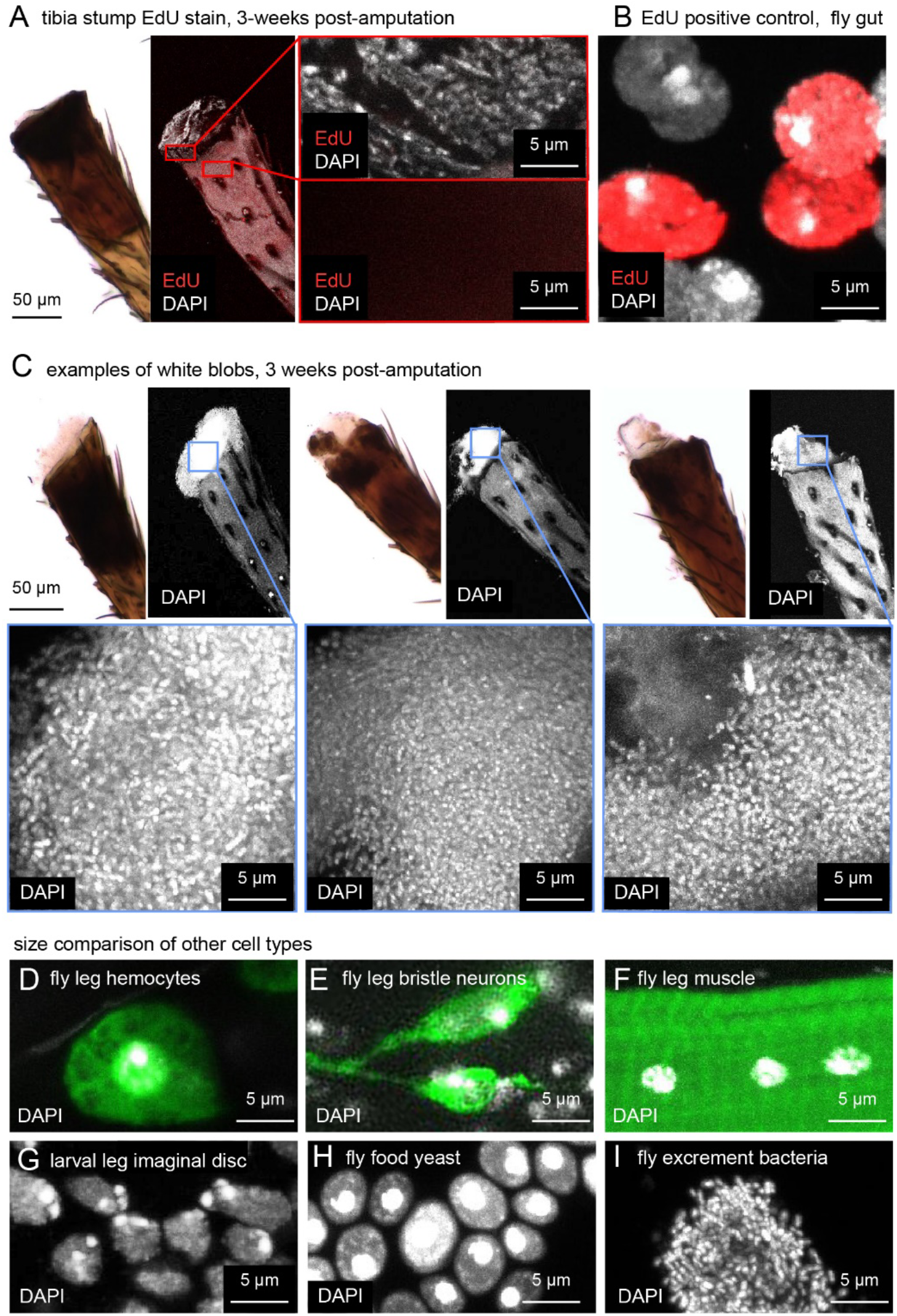
Investigation of the white blob on amputated tibia stumps. (A) Bright-field (left) and confocal images (right) of leg stumps stained with EdU (red) and DAPI (white) to test for cell proliferation. Tibia stumps did not incorporate EdU. The positive control, fly gut (B) did stain for EdU. (C) Three additional examples of DAPI staining of white blobs. Note that cells in (C) are smaller and more densely packed than fly leg hemocytes (D, green = *Hml>GFP*), fly leg bristle sensory neurons (E, green = *39A11>GFP*), fly leg muscle cells (F, green = phalloidin), larval leg imaginal disc cells (G), or fly food yeast (H). The nuclei in (C) are consistent with small, densely packed bacteria, such as those observed in fly excrement (I).

### The white blob on amputated leg stumps is likely a colony of bacteria

We found that the white blob stained robustly with the nuclear label DAPI **(Figure 3A, C)**. However, the nuclei in the white blob were about one tenth the size of other cells in the fly leg, including leg hemocytes **(Figure 3D)**, leg sensory neurons **(Figure 3E)**, and leg muscle **(Figure 3F)**. This discrepancy made us doubt that the white blob consisted of *Drosophila* cells. Abrams *et al*. also performed DAPI staining of the white blob, but the image in their paper (Figure 4f) lacked a scale bar, making it difficult to determine the source of the nuclei.

In some animals that do regenerate amputated limbs, such as newts and axolotls, blastema cells in an amputated leg de-differentiate, undergoing morphological and transcriptional changes to become more like younger cells (Leigh et al., 2018; Tanaka et al., 2016). To address this possibility in the fly leg, we compared DAPI staining in the white blob to DAPI staining in leg precursor cells of the larval leg imaginal disc **(Figure 3G)**. The nuclei in the white blob were again approximately one tenth the size of those of leg imaginal disc cells. This size difference is inconsistent with the idea that the white blob consists of de-differentiating fly tissue.

Finally, we compared DAPI staining of the white blob to other cells present in a *Drosophila* food vial: baker’s yeast and bacteria. The size of baker’s yeast nuclei was comparable to that of fly and other eukaryotic cells — significantly larger than the nuclei in the white blob **(Figure 3H)**. However, the size and density of DAPI labeling of the white blob did resemble DAPI staining of bacteria from fly excrement **(Figure 3I)**. We conclude that the white blob is consistent with a colony of bacteria and is not an intermediate structure related to regeneration morphology.

## Discussion

Our results support the conclusion that fly legs do not regenerate after amputation. Three weeks after amputation, we found that the tibia is completely devoid of neurons, muscles, and other cell nuclei. The stump thus appears to be simply a tube of hollow cuticle. DAPI and EdU staining of the amputation site confirmed the complete absence of living or regenerating cells in the tibia. Our results were not different between the control group and flies fed supplemental insulin, leucine, and glutamine. Overall, our conclusions are consistent with dogma that adult holometabolous insects do not regenerate lost ectodermal structures (Fox et al., 2020; Repiso et al., 2011).

How do we reconcile our results with those of Abrams *et al*.? We speculate that their conclusions were based on inaccurate measurement techniques. Detecting a subtle phenotype that occurs in only 1% of treated flies would require an exceptional degree of measurement accuracy. We propose that Abrams *et al*. quantification of sub-mm changes in tibia cuticle length lacked this requisite precision. Even so, this explanation fails to account for the differences in limb length distributions when the authors’ tracked limb length over time (e.g., Figure 5f in Abrams *et al*.). Another contributing factor could be bias in their experimental measurements or analysis. The paper states that, “blind measurements were performed on one pair of control and treated datasets”, but it is not clear how or on what data blinded measurements were performed.

Instead of relying on potentially noisy measurements of limb length over weeks, we used genetic tools and fluorescent labels to search for neurons, muscles, and other regenerated cells in amputated limbs. We found no evidence of living or regenerated cells within the tibia stump. We question how leg cuticle could regrow, even in rare cases, if no detectable living cells remain in the amputated stump. We also question how ingested insulin, one of the key ingredients in the dietary supplement, could affect limb regeneration, because it would be broken down in the gut (Winter, 1923). (Although it is possible it could access the amputated limb directly by physical contact between the stump and the food.) Finally, our evidence supports the conclusion that the blob of white tissue on the distal tip of the amputated tibia, which Abrams *et al*. claim is an intermediate regeneration morphology, is more likely a growth of bacteria.

Our new experimental results cast serious doubt on Abrams *et al*. conclusion that a diet supplement induces tibia regeneration in adult *Drosophila*. We did not attempt to reproduce or validate their regeneration results in jellyfish or mice, due to our lack of expertise with these species. Nonetheless, we feel that the flaws in execution and interpretation of the *Drosophila* experiments fundamentally undermine the sweeping conclusions of their paper. For example, they conclude, “that an inherent ability for appendage regeneration is retained in non-regenerating animals and can be unlocked with a conserved strategy.” Our results, and the past literature on regeneration in holometabolous insects, do not support this conclusion.

Our motivation to establish the truth about fly leg regeneration is more than academic. Promising experimental results in genetic model organisms, like flies and mice, often motivate experiments in other species, including humans. Because the ingredients in the supplemental diets used by Abrams *et al*. are already FDA approved, it is conceivable that “regeneration supplements” could be sold without rigorous prior testing. Based on the results in their paper, the authors have applied for a patent on, “Compositions and methods for inducing appendage and limb regeneration”. We feel that their data, and the efficacy of dietary supplements to induce regeneration, require additional scrutiny and independent replication.

## Materials and methods

| Table of genotypes |  |
| --- | --- |
| Figure 1, Figure 2B-D, Figure 3A-C, F-G | Wild type *Drosophila melanogaster* (Canton-S) |
| Figure 2A | w[*]; Mi{Trojan-GAL4.0}ChAT[MI04508-TG4.0] CG7715[MI04508-TG4.0-X]/ P{pJFRC7-020XUAS-IVS-mCD8::GFP}attP2 |
| Figure 3D | w[1118]; P{w[+mC]=Hml-GAL4.Delta}2, P{w[+mC]=UAS-2xEGFP}AH2 |
| Figure 3E | w[1118]; P{pJFRC7-020XUAS-IVS-mCD8::GFP}attP40; P{y[+t7.7] w[+mC]=GMR39A11-GAL4}attP2 |

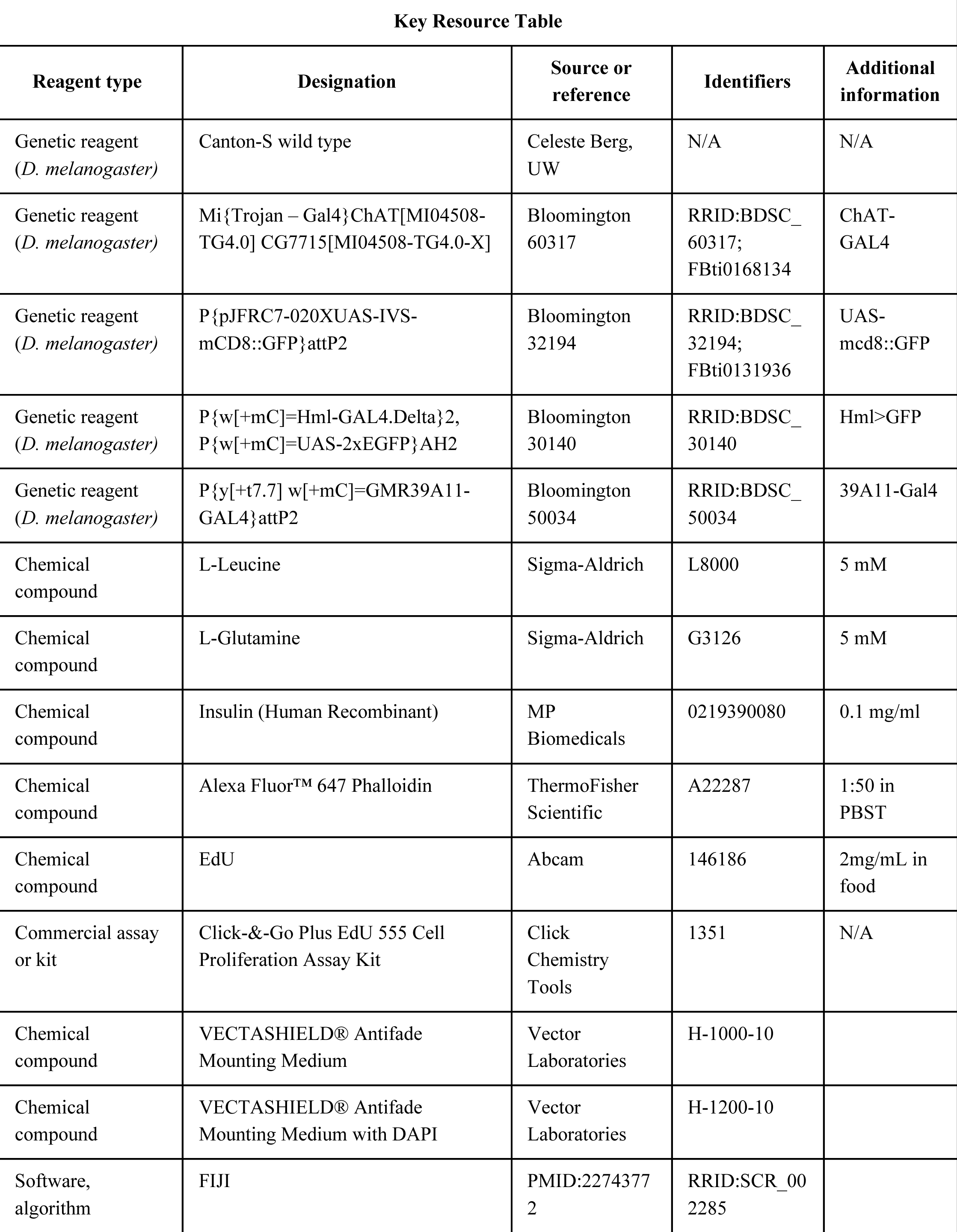

### Amputation and diet

*Drosophila* were raised on a standard cornmeal-molasses-yeast food fly food at 25°C with a 14 hr dark/10 hr light cycle. We used male and female adults, 1-2 days post-eclosion, reasoning that young flies would be more likely to regenerate and more likely to survive the three-week recovery period than old flies. For leg amputation, flies were anesthetized in groups of 20 on CO_2_ plates for 5 minutes or less. One hind-leg per fly was amputated at the mid-point of the tibia with a fine double-edge super-stainless razor blade (ASR 72-003). Amputated flies were included in our analysis only if the amputation site was within ∼50 μm of the tibia midpoint, using leg bristles as fiducial markers. Regeneration of cuticle was assessed according to whether the tibia length three weeks later fell outside of that range. After amputation, flies were immediately returned to a vial with either standard lab food or treated food, with random assignment.

To make treated food, vials of standard fly food were microwaved to liquefy the food. Before adding supplements, we let it cool to lukewarm to prevent the insulin from denaturing (Kaufmann et al., 2021). We added supplements in an aqueous stock solution and mixed the food for a final homogeneous concentration of 5 mM L-Leucine, 5 mM L-Glutamine, and 0.1 mg/ml insulin (Abrams et al., 2021). Food was mixed and allowed to set at room temperature for one hour. Flies were moved onto freshly prepared food every 2-3 days.

### Fixing, staining, and analysis

Legs or imaginal discs were fixed in 4% formaldehyde (PFA) PBS solution for 20 min followed by rinsing in PBS with 0.2% Triton X-100 (PBT) three times. To label muscle, legs were incubated in 1:50 phalloidin in a PBS solution with the following reagents to improve tissue penetrance: 1% triton X-100, 0.5% DMSO, 0.05 mg/ml Escin (Sigma-Aldrich, E1378), and 3% normal goat serum. Legs were allowed to incubate for one week at 4°C with occasional rocking. After staining, legs were rinsed 3x with PBS-Tx, 1x with PBS, and mounted onto slides in Vectashield with or without DAPI.

Each slide was labeled according to experimental condition. Prior to analysis, we taped-over the labels. Categorizations in Table 1 were performed with the experimenter blinded to experimental condition.

### DAPI staining of yeast and fly excrement bacteria

Cells were transferred to a slide, diluted in water, flame-fixed, then mounted in Vectashield with DAPI.

### EdU labeling

We supplemented the food recipe above with 2mg/mL EdU. Flies were moved to freshly prepared food every 2-3 days. After three weeks, legs and guts (positive control) were dissected and fixed in 4% formaldehyde (PFA) PBS solution for 20 min and processed according to Click-&-Go kit instructions. After staining, legs were rinsed 3x with PBS-Tx, 1x with PBS, and mounted in Vectashield with DAPI.

### Imaging

Mounted legs were imaged on a Confocal Olympus FV1000 (phalloidin, ChAT, EdU, and cuticle autofluorescence images) Leica DMI6000 Widefield (brightfield images), and Leica SP8X (DAPI images). Image stacks were processed in FIJI(Schindelin et al., 2012). Bright-field images were processed in Photoshop with the color channel mixer to correct a bluish background to truer white background.

## Acknowledgments

We thank Richard Mann, Michael Dickinson, Ben Ewen-Campen, and Bing Brunton for comments on the manuscript. We thank Beth Traxler for advice on identifying bacteria. We thank Rachel Wong for confocal use. We used stocks obtained from the Bloomington Drosophila Stock Center (NIH P40OD018537). We acknowledge support from the NIH (S10 OD016240) to the Keck Imaging Center at UW, and the assistance of its manager, Nathaniel Peters. This work was supported by a Pew Biomedical Scholar Award, a McKnight Scholar Award, the New York Stem Cell Foundation, and NIH grants R01NS102333 and U19NS104655 to JCT. JCT is a New York Stem Cell Foundation – Robertson Investigator.

## Notes

### Competing Interest Statement

The authors have declared no competing interest.

